# Aberrant localization of CDC42 C-terminal variants to the Golgi apparatus drives pyrin inflammasome-dependent autoinflammation

**DOI:** 10.1101/2021.09.03.458902

**Authors:** Masahiko Nishitani-Isa, Kojiro Mukai, Yoshitaka Honda, Hiroshi Nihira, Takayuki Tanaka, Hirofumi Shibata, Kumi Kodama, Eitaro Hiejima, Kazushi Izawa, Yuri Kawasaki, Mitsujiro Osawa, Yu Katata, Sachiko Onodera, Tatsuya Watanabe, Shigeo Kure, Junko Takita, Osamu Ohara, Megumu K. Saito, Ryuta Nishikomori, Tomohiko Taguchi, Yoji Sasahara, Takahiro Yasumi

**Author notes:** These authors contributed equally to this work (Masahiko Nishitani-Isa, Kojiro Mukai, Yoji Sasahara). Corresponding authors: Takahiro Yasumi and Yoji Sasahara, Takahiro Yasumi, Department of Pediatrics, Kyoto University Graduate School of Medicine, 54 Shogoin-Kawahara-cho, Sakyo-ku, Kyoto 606-8507, JAPAN, Yoji Sasahara, Department of Pediatrics, Tohoku University Graduate School of Medicine, 1-1 Seiryo-machi, Aoba-ku, Sendai, 980-8574, JAPAN.

## Abstract

Mutations in the C-terminal region of the *CDC42* gene cause severe neonatal-onset autoinflammation. Elevated levels of serum IL-18 in patients and effectiveness of IL-1β-blocking therapy indicate that the pathology involves abnormal inflammasome activation; however, the mechanism underlying autoinflammation remains to be elucidated. Using induced-pluripotent stem cells established from patients carrying *CDC42*^*R186C*^, we found that patient-derived cells secreted larger amounts of IL-1β in response to pyrin-activating stimuli. Aberrant palmitoylation and localization of CDC42^R186C^ protein to the Golgi apparatus promoted pyrin inflammasome assembly downstream of pyrin dephosphorylation. Aberrant subcellular localization was the common pathological feature shared by CDC42 C-terminal variants with inflammatory phenotypes, including CDC42^*192C*24^ that also localizes to the Golgi apparatus. Furthermore, the level of pyrin inflammasome overactivation paralleled that of mutant protein accumulation in the Golgi apparatus, but no that of the mutant GTPase activity. These results reveal an unexpected association between CDC42 subcellular localization and pyrin inflammasome activation that could pave way for elucidating the mechanism of pyrin inflammasome formation.

## Introduction

Autoinflammatory disorders are caused by dysregulated activation of innate immune systems and typically present in early childhood with fever and disease-specific patterns of organ inflammation. The concept of autoinflammatory diseases was proposed in 1999, and since that time a growing number of genetic causes have been identified for a variety of diseases (1, 2). Disease-based research has revealed the molecular mechanisms of excessive innate immune responses that drive autoinflammatory phenotypes, and these discoveries have provided us with novel therapeutic targets that might be used to effectively treat these conditions (3).

C-terminal variants of cell division control protein 42 homolog (CDC42) were recently shown to cause severe inflammation that presents in the neonatal period with fever, rashes, and a significant increase in inflammatory markers (4–6). Among the three reported variants (c.556C>T, *CDC42*^*R186C*^; c.563G>A, *CDC42*^*C188Y*^; c.576A>C, *CDC42*^*\*192C*24*^), *CDC42*^*R186C*^ induces the most severe phenotype. A unique property of CDC42^R186C^ is its aberrant localization to the Golgi apparatus, and previous studies suggested that impaired NK cell function or enhanced NF-κB signaling are associated with inflammation in the patients (5, 6). However, elevation of serum IL-18 levels and the effectiveness of IL-1β blocking reagents indicate that activation of an inflammasome is central to pathogenesis.

Studies of autoinflammatory pathology in humans have often been hampered by limited availability of patient samples. To overcome this difficulty, we used induced-pluripotent stem cells (iPSCs) established from the patients (7). iPSC-derived myeloid cell lines (iPS-MLs) can be obtained by introducing three transgenes into iPSC-derived monocytes (8). iPS-MLs phenotypically and functionally resemble primary immature macrophages and can be expanded in the presence of M-CSF. iPS-MLs can fully differentiate into macrophages (iPS-MPs), which produce higher levels of IL-1β and IL-18. These cell lines are useful for studying human autoinflammatory disorders, especially for analyses of the pyrin inflammasome. By contrast, it is difficult to establish a mouse disease model because murine pyrin lacks the B30.2/SPRY domain, where most mutations associated with familial Mediterranean fever (FMF) cluster (9).

In this study, we established iPS-MLs and iPS-MPs from patients carrying the *CDC42*^R186C^ mutation and demonstrated that the pyrin inflammasome plays an important role in driving hyperinflammation.

## Results

### CDC42^R186C^ triggers aberrant activation of pyrin inflammasome

Continuous elevation of IL-18 in the sera of the two patients with the *CDC42*^*R186C*^ variant (Supplemental Figure 1 and Case Report), along with previous reports describing a therapeutic benefit of anti-IL-1β treatment (4, 5), suggested that inflammasome activation is central to the pathogenesis of *CDC42*-associated autoinflammation. To determine whether a specific inflammasome is overactivated in patient cells, we established iPS-MLs and iPS-MPs and evaluated their response to various inflammasome activators. Compared with WT counterparts, iPS-MPs derived from patient 1 (Pt.1) produced higher levels of IL-1β and IL-18 in response to *Clostridium difficile* Toxin A (TcdA), a pyrin inflammasome activator, but not other inflammasome activators such as nigericin and flagellin (Figure 1A). iPS-MLs exhibited the same pattern, although the levels of IL-1β production were lower than in iPS-MLs, and IL-18 was undetectable (Figure 1B). Because IL-6 secretion did not differ between WT and patient-derived clones (Figure 1, A and B), the elevated production of IL-1β and IL-18 was likely due to an exaggerated response to TcdA rather than a difference in the induction efficacy of iPS-MPs or the genetic background of the cells. The same phenotype was confirmed in cells established from fibroblasts derived from Pt.2 (Supplemental Figure 2, A and B). Enhanced ASC speck formation in patient-derived iPS-MPs was evident after TcdA stimulation (Figure 1C).

**Figure 1.**
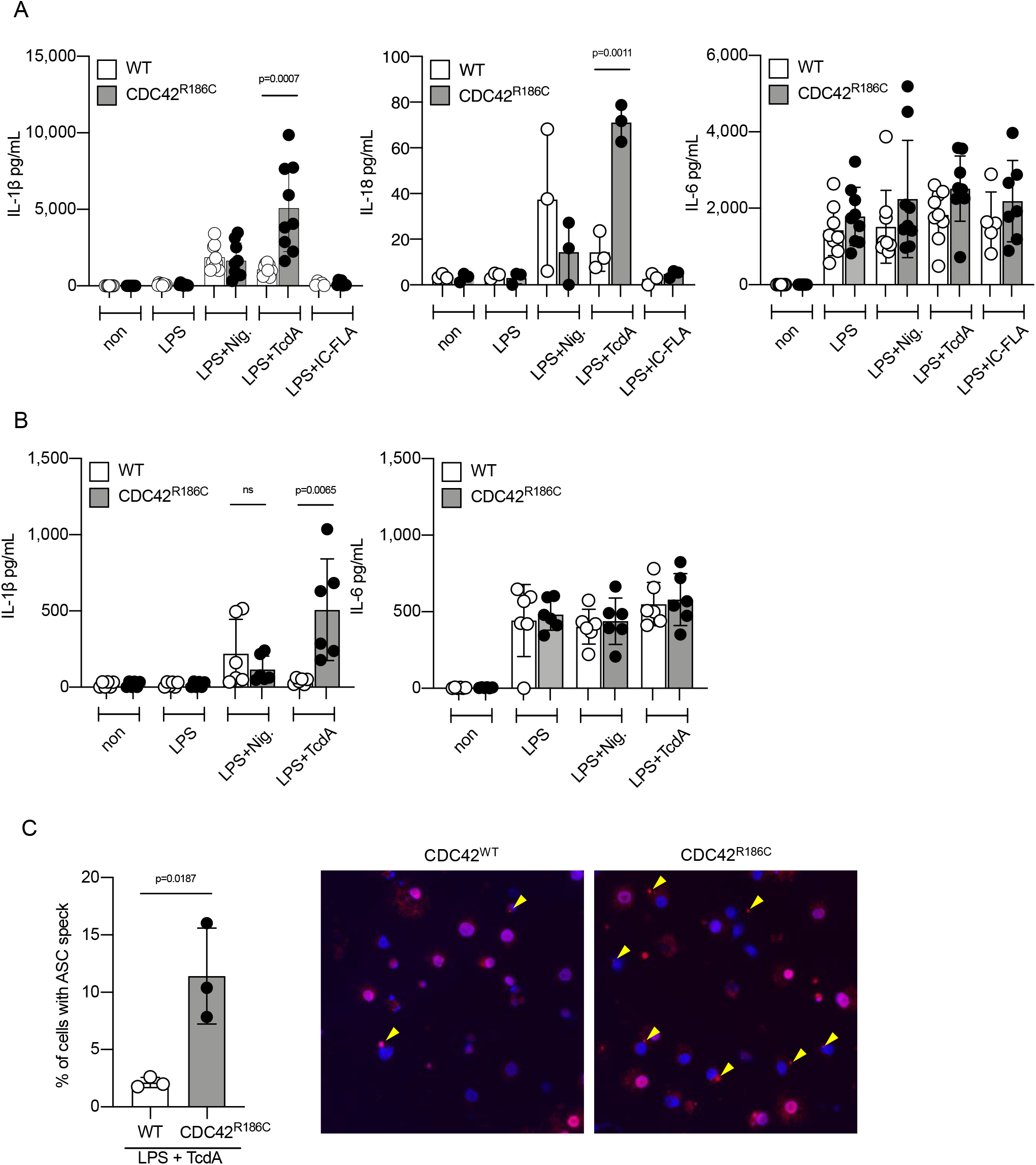
Response to TcdA stimulation is elevated in iPS-MPs and iPS-MLs derived from a patient carrying CDC42^R186C^. IL-1β, IL-18, and IL-6 release in response to various inflammasome stimuli from iPS-derived (**A**) MPs and (**B**) MLs established from Pt.1 and healthy controls. The results are from multiple experiments using three independent iPS clones. (**C**) Ratio and representative images of ASC speck formation (arrowheads) in iPS-MPs from Pt.1 and healthy controls after priming with LPS and TcdA. The results are representative of three independent experiments with similar results. Statistical significance was determined by one-way ANOVA with Tukey’s multiple comparison test in (**A**) and (**B**), and by unpaired t-test in (**C**).

Increased IL-1β production from *CDC42*^*R186C*^ iPS-MLs was inhibited by pyrin inflammasome inhibitors such as colchicine, ABT-751, and CA4P, but not by a NLRP3 inflammasome inhibitor MCC 950 (Figure 2A). IL-1β overproduction and enhanced ASC speck formation in patient-derived clones were suppressed by siRNA-mediated silencing of the *MEFV* gene with no obvious effect on IL-6 production (Figure 2, B-D). Furthermore, single-base genomic correction of *CDC42*^R186C^ abolished IL-1β overproduction upon LPS and TcdA stimulation while IL-6 production levels remained unchanged (Figure 2E). These findings all confirmed that *CDC42*^R186C^ led to hyperactivation of pyrin inflammasome.

**Figure 2.**
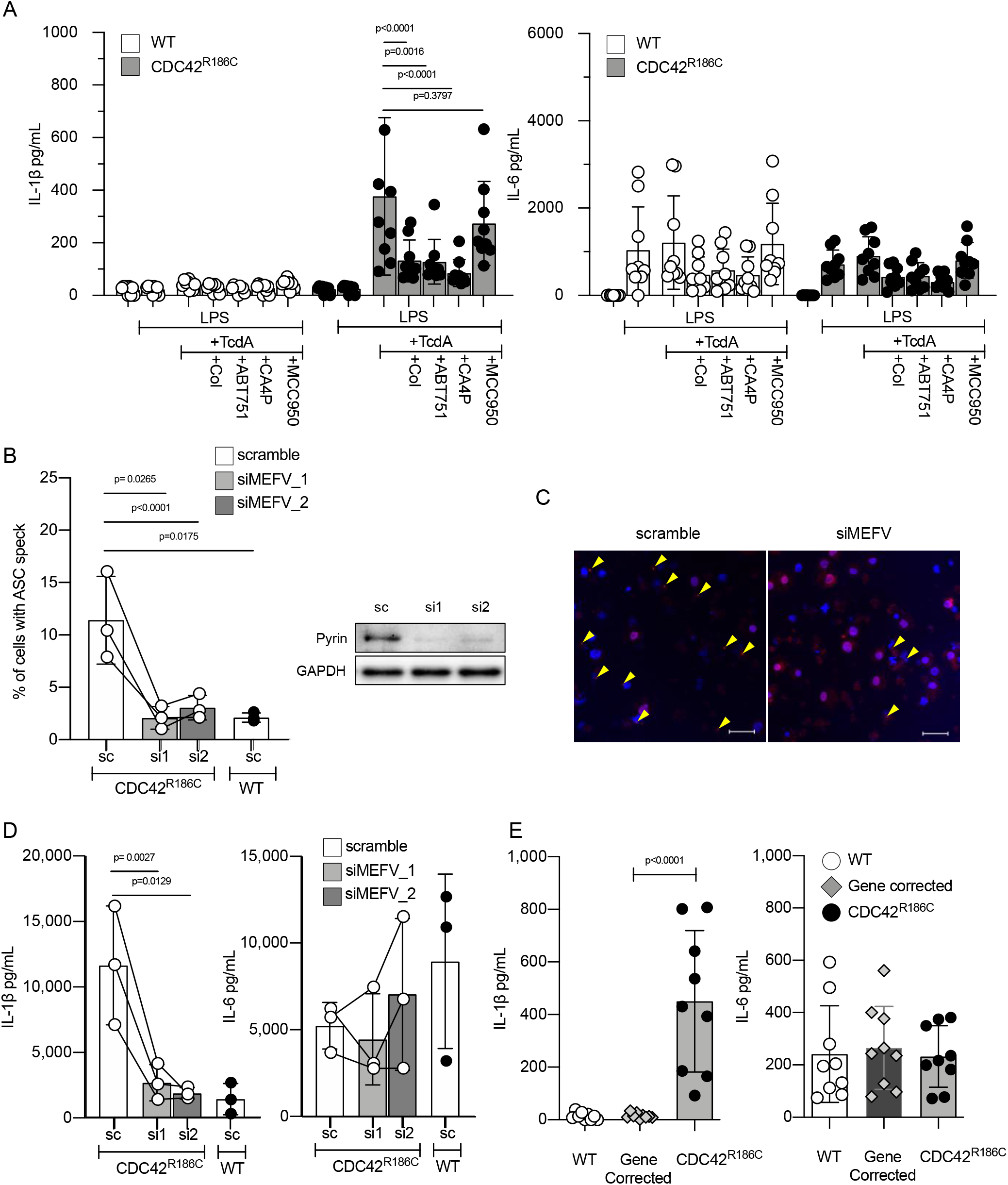
CDC42^R186C^ induces pyrin inflammasome hyperactivation. (**A**) IL-1β and IL-6 release from iPS-MLs derived from Pt.1 and healthy controls stimulated with LPS and TcdA in the presence or absence of pyrin and NLRP3 inhibitors. (**B**) Ratio and (**C**) images of ASC speck formation, and (**D**) release of IL-1β and IL-6 from Pt.1-derived iPS-MPs in which *MEFV* was knocked down by siRNA. Immunoblot images in (**B**) show the effects of siRNAs. (**E**) IL-1β and IL-6 release from Pt.1-derived iPS-MLs, mutation-corrected Pt.1-derived iPS-MLs, and iPS-MLs from healthy controls stimulated with LPS+TcdA. The results are representative of three independent experiments with similar results. Statistical significance was determined by one-way ANOVA with Tukey’s multiple comparison test in (**A**), and by ratio-paired t test in (**B**), (**D**), and (**E**).

### Aberrant palmitoylation causes trapping of CDC42^R186C^ protein in the Golgi apparatus and induces pyrin activation

Heterozygous mutations of the *CDC42* gene are responsible for Takenouchi-Kosaki syndrome (TKS), which is characterized by developmental delay, facial dysmorphism, and macrothrombocytopenia (10–12). However, inflammatory manifestations are exceptional in TKS, whereas our patients presented with lethal inflammation soon after birth with no signs of facial dysmorphism or abnormal platelet size. These all suggest that *CDC42*^*R186C*^ induces different pathophysiology than TKS-associated mutations.

To explore the mechanism of *CDC42*^R186C^-induced aberrant activation of the pyrin inflammasome, we first compared the GTPase activity of various *CDC42* mutants in HEK293T cells by pull-down assay. As previously reported (12), TKS-associated *CDC42* mutants exhibited diverse derangement in their GTPase activities (Figure 3A). However, *CDC42*^*R186C*^ did not exhibit a significant change in its GTPase activity, consistent with the fact that this mutation is far from the GTP binding site (Figure 3A). Instead, the expression level of CDC42^R186C^ protein was elevated, and the electrophoresis mobility was slightly altered (Figure 3A and Supplemental Figure 4). This prompted us to check the subcellular distribution of CDC42^R186C^ protein. Consistent with the previous reports, we found that the protein was retained in the Golgi apparatus (5, 6), a striking feature that is not observed in TKS-associated mutations (Figure 4).

**Figure 3.**
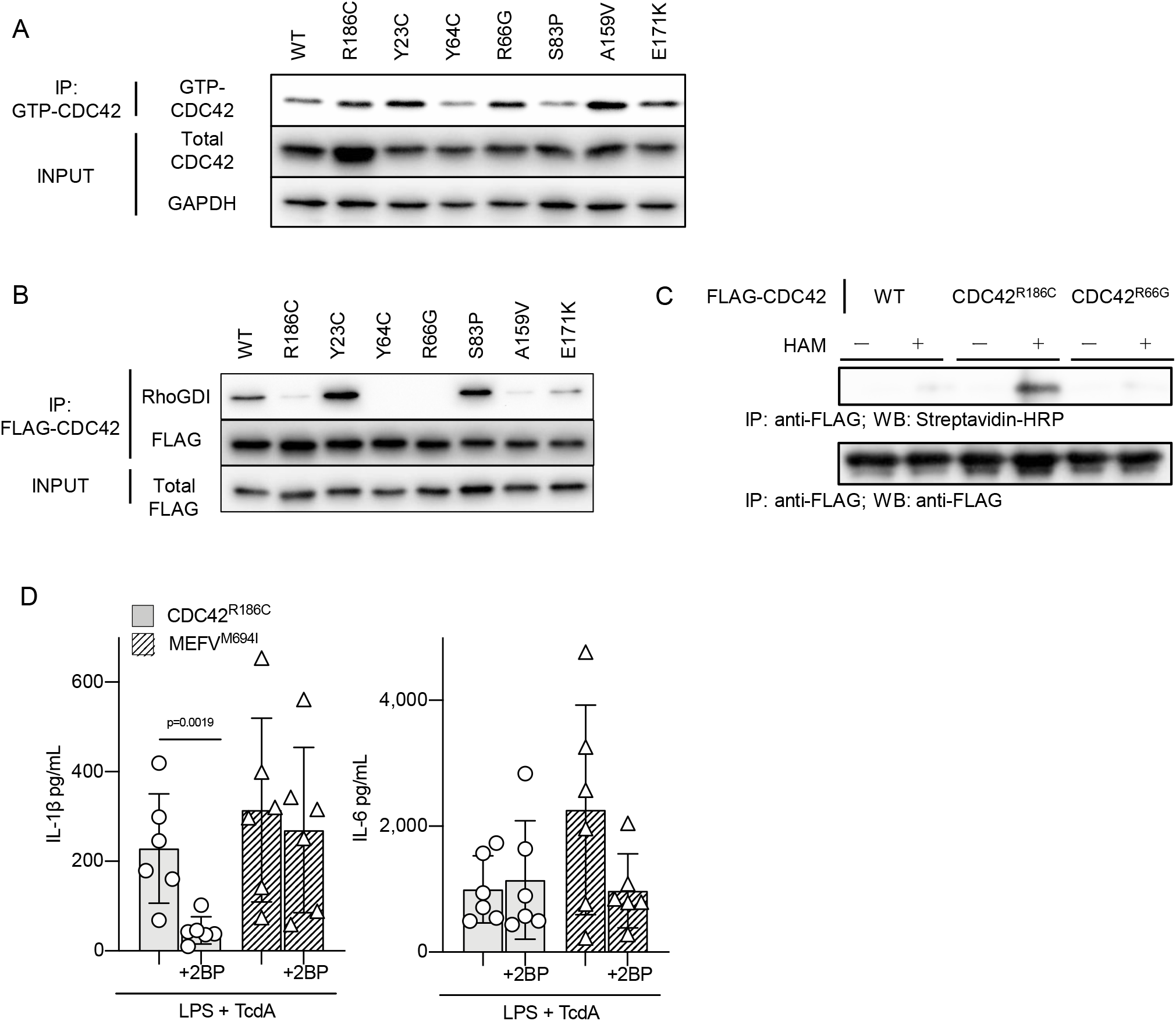
Palmitoylation of CDC42^R186C^ variant causes IL-1β overproduction. (**A)** Immunoblot of GTP-bound CDC42 and total CDC42 in the lysates of HEK293T cells transiently expressing CDC42 variants. (**B)** Interaction of CDC42 variants with Rho-GDI assessed by immunoprecipitation (IPs) of FLAG-tagged CDC42 transiently expressed in HEK293T cells. (**C)** Acyl–biotin exchange assay to evaluate S-palmitoylation levels of FLAG-CDC42 variants expressed in HEK293T cells with or without hydroxylamine (HAM) treatment. (**D)** IL-1β and IL-6 release from iPS-MPs carrying CDC42^R186C^ and MEFV^M694I^ cultured overnight with or without 2-bromo-palmitate (2BP). Representative results of three independent experiments are shown for **A** to **C**, and the results in **D** are from two independent experiments with three iPS clones. Statistical significance was determined by one-way ANOVA with Tukey’s multiple comparison test in **D**.

**Figure 4.**
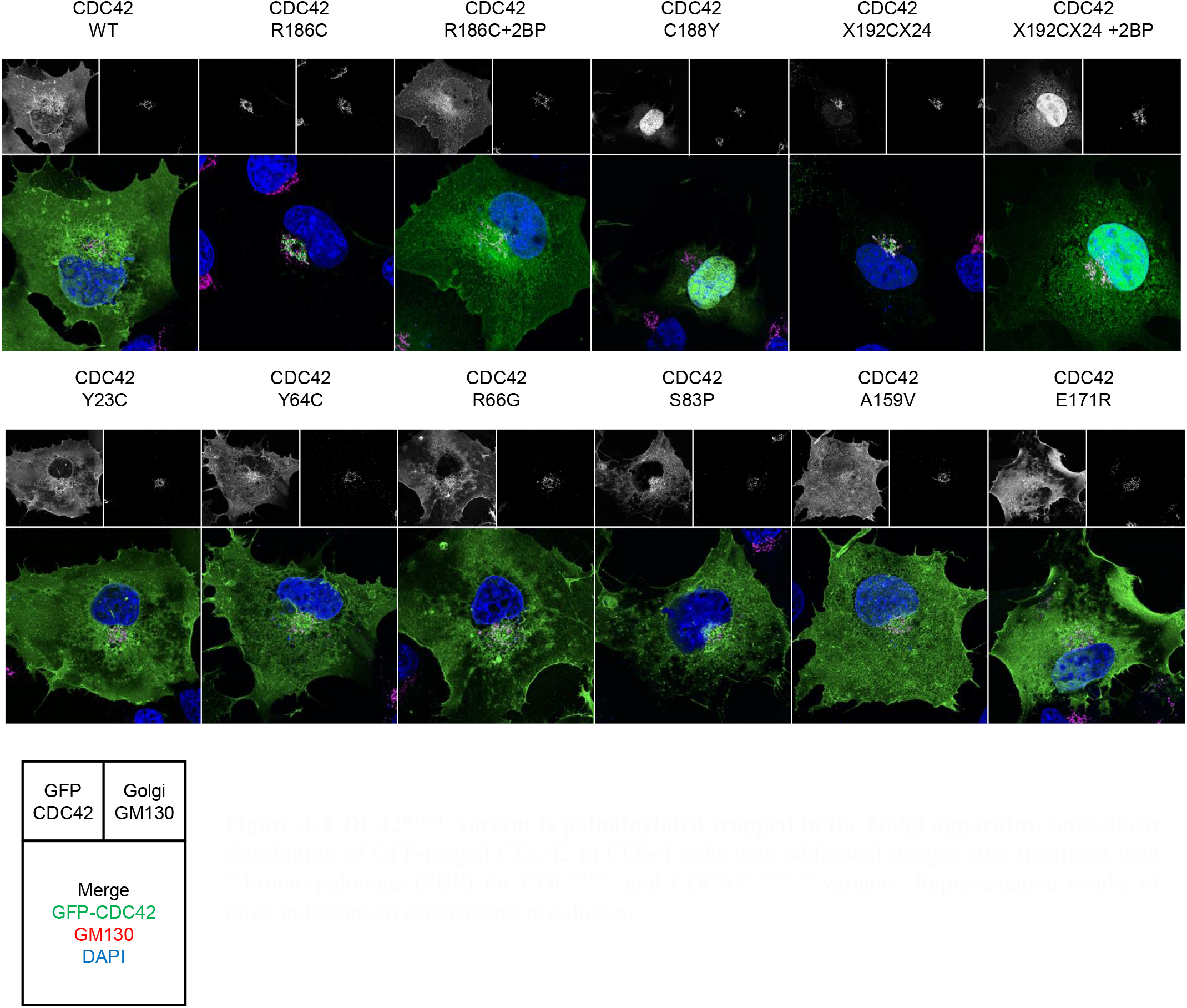
CDC42^R186C^ variant is palmitoylated trapped in the Golgi apparatus. Subcellular distribution of GFP-tagged CDC42 in COS-1 cells with additional images after treatment with 2-bromo-palmitate (2BP) for CDC^R186C^ and CDC42^*192C*24^ variants. Representative results of three independent experiments are shown.

To determine the molecular basis for the abnormal localization of CDC42^R186C^ protein to the Golgi apparatus, we assessed the binding of each CDC42 mutant to Rho GDP dissociation inhibitors (GDIs), which modulate the subcellular localization of CDC42. Binding of CDC42^R186C^ protein to Rho-GDI was impaired, as was the binding of some TKS mutants not associated with an inflammatory phenotype, such as CDC42^R66G^ (Figure 3B); however, the subcellular distributions of the TKS mutants were comparable to that of WT CDC42 protein (Figure 4). *CDC42*^*R66G*^ iPS-MP clones generated by single-base editing did not produce higher levels of IL-1β in response to LPS and TcdA stimulation (Supplemental Figure 3). These results suggest that impaired binding of CDC42^R186C^ protein to Rho-GDI is not critical for its localization to the Golgi apparatus or to pyrin inflammasome hyperactivation.

Bekhouche, et al. showed that abnormal palmitoylation of CDC42^R186C^ causes its abnormal localization to Golgi apparatus (6). The *CDC42* transcript is alternatively spliced to produce two isoforms. CDC42 isoform 1 is ubiquitously expressed, and its C-terminal region encodes a -CAAX motif that is geranylgeranylated at Cys^188^, allowing it to be tethered to cell membranes (13). CDC42 isoform 2 is expressed in the central nervous system, and its C-terminal region is palmitoylated at Cys^186^, as in the case of the small GTPase H-Ras (14). Palmitoylation is a key regulatory step that determines the subcellular distribution of proteins, and the *CDC42*^*R186C*^ mutation creates a potential palmitoylation site, Cys^186^, at the C-terminal region of CDC42 isoform 1. By performing acyl–biotin exchange palmitoyl-protein purification from HEK293T cells transfected with CDC42 mutants, we confirmed that CDC42^R186C^, but not CDC42^WT^ or CDC42^R66G^, was palmitoylated (Figure 3C). We then assessed whether aberrant palmitoylation was responsible for the retention of CDC42^R186C^ protein to the Golgi apparatus in HEK293T cells, and for IL-1β overproduction by CDC42^R186C^ iPS-MLs, by treating cells with the general protein palmitoylation inhibitor 2-bromo-palmitate (2BP). 2BP treatment of HEK293T cells transfected with the *CDC42*^*R186C*^ mutant decreased its retention in the Golgi apparatus (Figure 4). Furthermore, 2BP treatment significantly suppressed overproduction of IL-1β from *CDC42*^*R186C*^ iPS-MLs, but not from iPS-MLs derived from FMF patients carrying *MEFV*^*M694I*^ (Figure 3D). These data indicate that aberrant palmitoylation of CDC42^R186C^ protein caused its retention in the Golgi apparatus and triggered overactivation of the pyrin inflammasome in response to LPS and TcdA stimulation.

### CDC42^R186C^ enhances pyrin inflammasome assembly

We then tried to unravel the molecular mechanism of pyrin inflammasome overactivation in *CDC42*^R186C^ cells. Silencing of *CDC42* transcripts suppressed IL-1β production in both WT and *CDC42*^R186C^ iPS-MPs after stimulation with TcdA (Figure 5A). Moreover, it also suppressed IL-1β production from these cells after nigericin stimulation (Figure 5A), suggesting that CDC42^WT^ is required for full activation of both pyrin and NLRP3 inflammasomes. Together, these observations indicate that CDC42^R186C^ is not a simple gain-of-function variant, as it selectively enhanced pyrin but not NLRP3 inflammasome activation.

**Figure 5.**
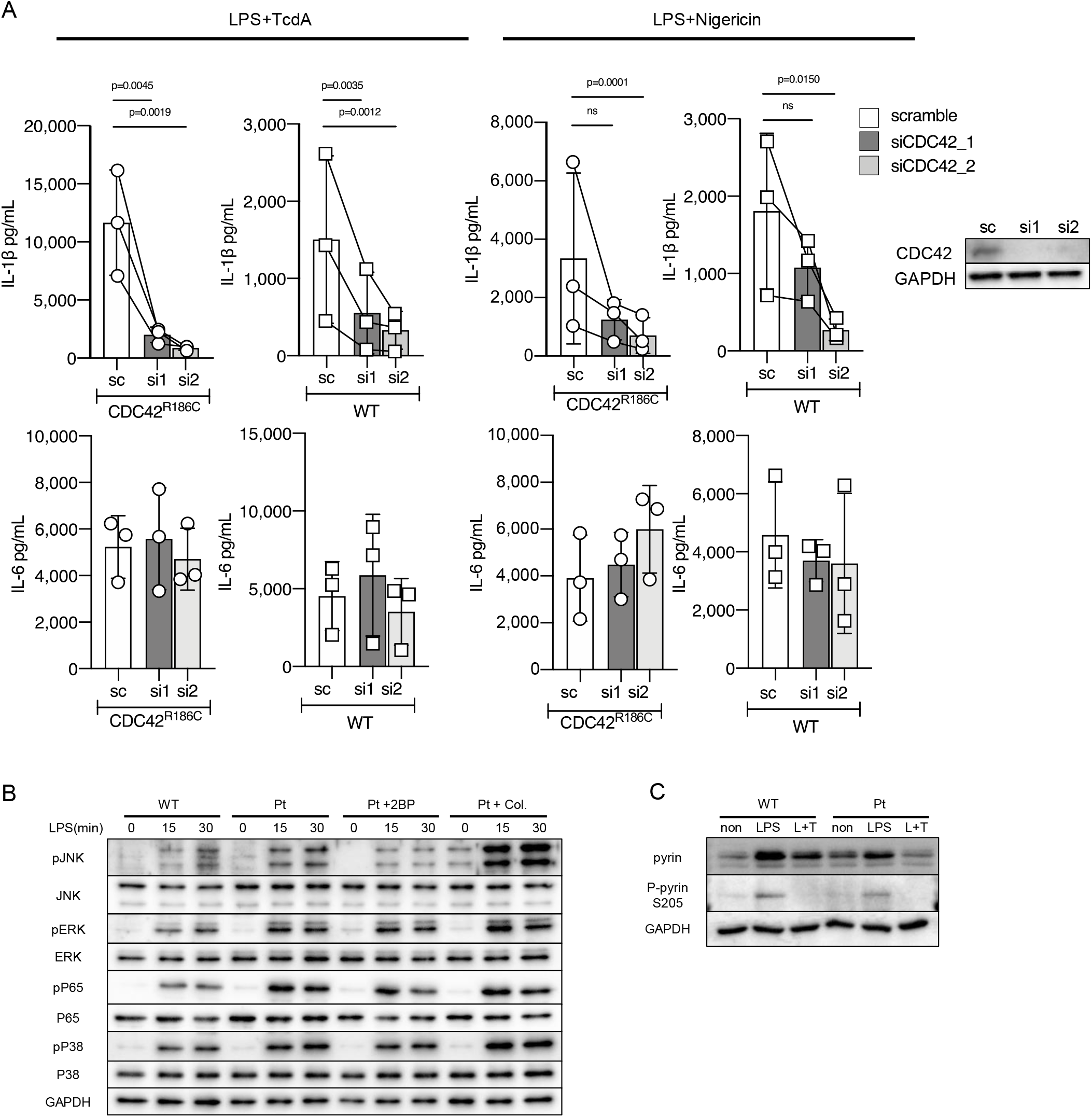
CDC42^WT^ is required for full activation of pyrin and NLRP3 inflammasomes, and NF-kB signaling is elevated in patient-derived iPS-MLs. (**A**) iPS-MLs from Pt.1 and healthy controls were treated with scramble or *CDC42* siRNA and the release of IL-1b and IL-6 was monitored in response to LPS+TcdA or LPS+ nigericin stimulation. Immunoblot images show the effects of siRNAs. A representative result of three independent experiments with three clones is shown. Immunoblot of (**B**) JNK, ERK, P65, P38, and (**C**) pyrin molecules and their phosphorylated forms in iPS-MLs from Pt.1 and a healthy control in response to LPS and TcdA stimuli. Representative results of three independent experiments with similar results are shown.

We next investigated the involvement of the NF-κB pathway, as a previous study suggested that elevated signaling is associated with the inflammatory phenotype of the patients (6). By analyzing the signaling pathways downstream of LPS priming, we found that phosphorylation of JNK, ERK, p65, and p38 was elevated in *CDC42*^*R186C*^ cells (Figure 5B). However, 2BP treatment abolished phosphorylation of JNK, but not that of other molecules; indeed, colchicine augmented phosphorylation of the other molecules (Figure 5B). Furthermore, pyrin expression did not differ between WT and the mutant cells after LPS priming (Figure 5C). These results rule out a pivotal role of the MAPK and NF-κB pathways in pyrin inflammasome overactivation.

To further determine how the *CDC42*^R186C^ mutant selectively modulates pyrin inflammasome activation, we compared the characteristics of cells carrying *CDC42*^R186C^ and *MEFV*^M694I^ mutations. Pyrin is phosphorylated at Ser^208^ and Ser^242^ by protein kinase N1 and N2 (PKN1/2), members of the protein kinase C (PKC) family, and at steady state it is kept inactive by inhibitory 14-3-3 proteins (15–18). Recent evidence suggests that the pyrin inflammasome is controlled by two independent mechanisms in healthy donors: pyrin dephosphorylation by inhibition of PKN1/2, followed by inflammasome maturation involving microtubule dynamics. Pyrin molecules harboring FMF-associated *MEFV* mutations, such as *MEFV*^M694I^, are vulnerable to dephosphorylation stimuli (19). Overproduction of IL-1β from both *CDC42*^R186C^ and *MEFV*^M694I^ iPS-MLs was inhibited to similar extents by bryostatin (Figure 6A), an activator of PKC that phosphorylates pyrin. The inhibitory effect of colchicine, which works predominantly by altering microtubule dynamics, was more potent in *MEFV*^M694I^ iPS-MLs than in *CDC42*^R186C^ iPS-MLs (Figure 6B). In addition, stimulation with UCN01, which dephosphorylates pyrin, induced higher levels of IL-1β production in *CDC42*^R186C^ iPS-MLs than in *MEFV*^M694I^ iPS-MLs (Figure 6C). Together, these results suggest that CDC42^R186C^ mutant promotes microtubule-dependent assembly of the pyrin inflammasome once pyrin is dephosphorylated.

**Figure 6.**
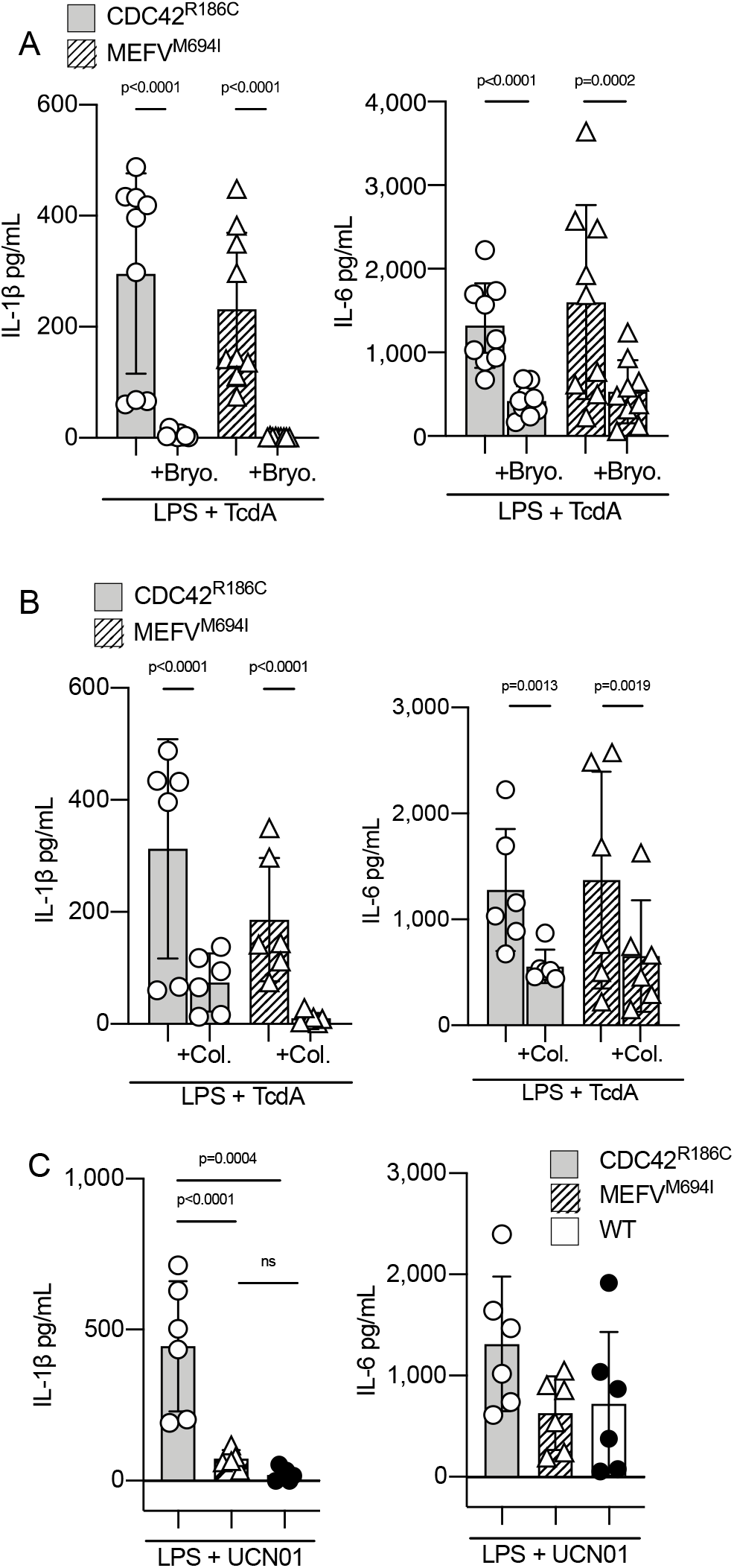
CDC42^R186C^ promotes pyrin inflammasome assembly. Effect of (**A**) bryostatin and (**B**) colchicine on the release of IL-1β and IL-6 from iPS-MLs derived from patients with CDC42^R186C^ and MEFV^M994I^ in response to LPS+TcdA stimulation. (**C**) IL-1β and IL-6 release from iPS-MLs derived from healthy donors and patients with CDC42^R186C^ and MEFV^M994I^ after stimulation with UCN01. Results are from two independent experiments using three independent clones. Statistical significance was determined by ratio-paired t test in (**A**) and (**B**), and by one-way ANOVA with Tukey’s multiple comparison test in (**C**).

### Aberrant subcellular distribution is the common finding of inflammatory CDC42 C-terminal variants

We then asked whether other inflammation-associated CDC42 C-terminal variants would have biomolecular characteristics similar to those of CDC42^R186C^. Indeed, we found that CDC42^*192C*24^ is trapped in the Golgi apparatus, like CDC42^R186C^ (Figure 4), although less of the former protein accumulated, likely reflecting a difference in the stability of CDC42^R186C^ and CDC42^*192C*24^ mutant proteins (Supplemental Figure 4). Furthermore, 2BP treatment abolished retention of CDC42^*192C*24^ in the Golgi apparatus (Figure 4), indicating that CDC42^*192C*24^ is also aberrantly palmitoylated. Surprisingly, CDC42^C188Y^ was predominantly distributed in the cytoplasm and nucleus, with little retention in the Golgi apparatus (Figure 4). These results clearly demonstrate that the inflammatory phenotype of patients is associated with aberrant subcellular distribution of CDC42 C-terminal variants.

### CDC42^*192C*24^ also induces pyrin inflammasome overactivation

Finally, we asked whether CDC42^C188Y^ and CDC42^*192C*24^ variants induce hyperactivation of the pyrin inflammasome similarly to CDC42^R186C^. To determine this, we engineered iPSCs to carry CDC42^WT^, CDC42^R186C^, CDC42^C188Y^, and CDC42^*192C*24^ variants in the same genetic background using homologous recombination and evaluated IL-1β production from iPS-MPs. Compared with iPS-MPs carrying CDC42^WT^, iPS-MPs carrying CDC42^R186C^ and CDC42^*192C*24^, but not CDC42^C188Y^, variants produced higher levels of IL-1β specifically in response to LPS and TcdA stimulation. CDC42^R186C^ induced higher levels of IL-1β than CDC42^*192C*24^ (Figure 7A), paralleling the severity of inflammation reported in the patients and the extent of mutant protein accumulation in the Golgi apparatus. Furthermore, 2BP treatment suppressed production of IL-1β, but not IL-6, from CDC42^R186C^ and CDC42^*192C*24^ iPS-MPs (Figure 7B). Next, we transfected CDC42 variants with or without WT-pyrin into THP-1 monocytic cells and assessed their effect on induction of cell death (Figure 7, C and D; and Supplemental Figure 5). In the absence of pyrin co-expression, CDC42 variants induced similar levels of THP-1 cell death (Figure 7C). In the presence of WT-pyrin, CDC42^R186C^ and CDC42^*192C*24^, but not CDC42^C188Y^, increased the frequency of cell death of THP-1 cells to a greater extent than CDC42^WT^ or the non-inflammatory TKS-associated variant CDC42^R66G^ (Figure 7C); again, CDC42^R186C^ induced higher levels of cell death than CDC42^*192C*24^. Surprisingly, pyrin-dependent cell death induced by the CDC42^R186C^ and CDC42^*192C*24^ mutants was independent of their GTPase activity because the variants harboring additional T17N mutation, which renders these variants defective in nucleotide binding (20, 21), induced comparable levels of cell death in THP-1 cells (Figure 7D).

**Figure 7.**
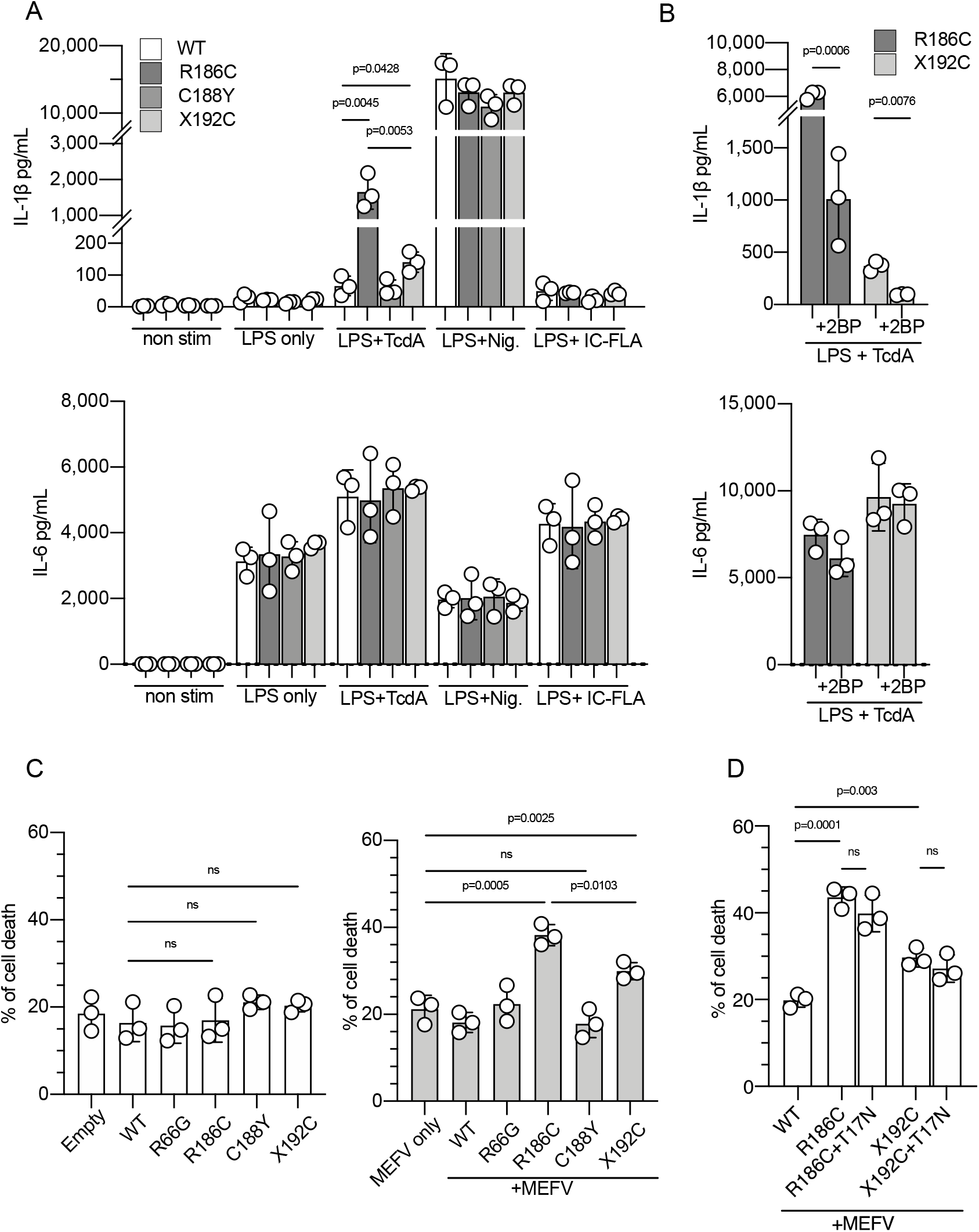
CDC42^*192C*24^ variant also induces elevated pyrin-inflammasome activation. (**A**) IL-1β and IL-6 release in response to various inflammasome stimuli from iPS-MPs harboring CDC42^WT^, CDC42 ^R186C^, CDC42^C188Y^, or CDC42^*192C*24^, all of which were generated from the same iPS clone; (**B**) the effect of 2BP in iPS-MPs harboring CDC42 ^R186C^ or CDC42^*192C*24^ is shown. Representative results of three independent experiments using three independent clones with similar results are shown. (**C**) THP-1 cells were nucleofected with WT or CDC42 variants alone or with pyrin, and the proportion of dead cells was evaluated 24 h later. (**D**) THP-1 cells were nucleofected with pyrin or CDC42 C-terminal variants including those with the additional T17N mutation, and the proportion of dead cells was evaluated 24 h later. Data obtained from three independent experiments are shown. Statistical significance was determined by one-way ANOVA with Tukey’s multiple comparison test.

Taken together, our results indicate that aberrant subcellular distribution of CDC42 C-terminal variants is associated with inflammatory phenotypes, and that palmitoylation-induced Golgi trapping of CDC42^R186C^ and CDC42^*192C*24^ variants drives pyrin-dependent autoinflammation.

## Discussion

CDC42, a member of the RAS superfamily of low-molecular-weight GTP/GDP-binding proteins, functions as a major node in intracellular signaling including adhesion, migration, polarity, and proliferation (22). CDC42 cycles between a GTP-bound active and a GDP-bound inactive state through regulation by GTPase-activating proteins and guanine nucleotide exchange factors (23). CDC42 has two isoforms. Isoform 1, which is ubiquitously expressed, is tethered to membranes through geranyl-geranylation at a C-terminal Cys^188^ (13). Isoform 2 is expressed in the brain and has alternative C-terminal amino acids, exchanging the prenylation motif for a sequence that is instead palmitoylated (14). The intracellular distribution of CDC42 is regulated by these C-terminal modifications and its binding affinity for Rho GDIs (24).

In 2015, heterozygous CDC42 mutations were reported to cause TKS, which is characterized by developmental delay, facial dysmorphism, and macrothrombocytopenia (10, 11). Martinelli et al. showed that GTPase activity and binding affinity for effector molecules varies greatly among TKS-associated CDC42 variants and have proposed phenotype–genotype classification of TKS into three groups (12); however, inflammatory manifestations are exceptional in TKS. In 2019, a study showed that heterozygous C-terminal variants of CDC42 isoform 1 cause severe and early-onset autoinflammation (4, 5). Notably in this regard, patients with the CDC42^R186C^ variant present with the most severe inflammation soon after birth but without signs of dysmorphism or macrothrombocytopenia, a phenotype consistent with our two cases (see Case Report). However, patients with other C-terminal variants of CDC42 isoform 1 present with both inflammatory and dysmorphic features (4). Recently, inflammation was also reported in a few patients with non–C-terminal CDC42 variants; however, in those individuals, the onset of inflammation was years to decades later, and the features of immunodeficiency were more prominent (25, 26). Together, these observations suggest that distinct and complex mechanisms underlie the inflammation and other clinical features caused by CDC42 variants.

In this study, we found that iPS-MLs and -MPs derived from patients carrying the CDC42^R186C^ variant formed higher levels of pyrin inflammasomes. As reported previously, a unique property of the CDC42^R186C^ mutant is its aberrant palmitoylation and localization to the Golgi apparatus (5, 6). Treatment with a palmitoyl transferase inhibitor released the mutant protein from the Golgi apparatus and normalized the secretion of IL-1β to WT levels. These data clearly illustrate that aberrant palmitoylation of CDC42^R186C^ plays a central role in the retention of the mutant to the Golgi apparatus, as well as in the exaggerated activation of the pyrin inflammasome.

Analysis of the mechanism of pyrin inflammasome overactivation by comparison with cells harboring *MEFV*^M694I^ revealed that the CDC42^R186C^ mutant likely promotes microtubule-dependent assembly of pyrin inflammasomes after pyrin dephosphorylation. Inflammatory pathology driven by the CDC42^R186C^ mutant could not be explained by alteration in GTPase activity or reduced binding affinity for GDIs, as some TKS-causative CDC42 variants that are not associated with inflammatory phenotypes exhibited more severe derangement of these properties. Rather, our results suggest that aberrant subcellular distribution of the variant protein is central to the inflammatory phenotype of the patients with CDC42 C-terminal variants. Due to aberrant palmitoylation, the CDC42^*192C*24^ mutant was also trapped in the Golgi apparatus and induced overactivation of the pyrin inflammasome. Notably, the level of IL-1β production paralleled the level of mutant protein accumulation in the Golgi apparatus. Furthermore, pyrin-dependent THP-1 cell-death induced by CDC42 C-terminal variants was independent of their GTPase activity. Strong association between the autoinflammatory phenotype and abnormal subcellular localization of the C-terminal CDC42 variants, but not with their altered GTPase activity or binding affinity for GDIs, indicates that ectopic expression of CDC42 molecule drives assembly of a specific inflammasome through non-physiological pathways.

We acknowledge that the study has several limitations. First, we were unable to reproduce the inflammatory phenotype of cells carrying the CDC42^C188Y^ variant. However, it is possible that the predominantly cytosolic distribution of this mutant is associated with abnormal activation of other inflammasomes. It is also possible that the CDC42^C188Y^ variant may induce inflammation only in specific cell types, such as granulocytes and tissue-resident macrophages, which have different characteristics than iPS-MLs and -MPs. Second, we could not elucidate the molecular mechanism of enhanced pyrin inflammasome assembly induced by Golgi-trapped CDC42 variants. Further studies are needed to unravel the mechanism that drives hyperactivation of pyrin and possibly other inflammasomes by C-terminal CDC42 variants.

Our results are consistent with a previous study that reported elevated NF-κB signaling in patient cells (6), although it is unlikely that this pathway plays a major role in inducing pyrin inflammasome hyperactivation. However, we believe that this activation of NF-κB signaling contributes to the inflammatory phenotype in patients, especially skin manifestations, as the patient described in that study continued to have skin inflammation even after receiving hematopoietic cell transplantation that resulted in complete donor chimerism (6). In regard to another important inflammatory phenotype seen in the patients, it is possible that CDC42^R186C^ impairs NK function and contributes to the development of hemophagocytic lymphohistiocytosis (HLH) (5). We analyzed NK cells and cytotoxic T lymphocytes (CTLs) derived from Pt.1, but found no significant disturbance in degranulation of both cell types or NK cell cytolytic activity (Supplemental Figure 5). Because high levels of serum IL-18 cause secondary impairment of NK cell function, which is associated with the development of macrophage activation syndrome (27, 28), we believe that the HLH phenotype seen in the patients is likely to represent a secondary form. However, because we used cells that were expanded and stimulated with IL-2, which can overcome mild derangement of cytolytic lymphocyte functions (29, 30), we cannot rule out the possibility that CDC42^R186C^ itself mildly impairs these functions.

Of clinical importance, our results suggest that colchicine and other microtubule inhibitors could be used to treat these patients. Because IL-1β blocking therapy was not sufficient to suppress inflammation, and only one patient carrying the CDC42^R186C^ variant has survived for a long period of time without receiving hematopoietic cell transplantation, it would be highly beneficial if these inhibitors were effective in suppressing inflammation.

In conclusion, we have shown that all inflammatory CDC42 C-terminal variants are aberrantly distributed subcellularly, and that palmitoylation-induced trapping of the mutant protein to the Golgi apparatus triggers overactivation of the pyrin inflammasome. Further studies are needed to unravel the complex involvement of CDC42 in the activation of inflammasomes; the resultant knowledge could lead to the development of better management strategies for the patients.

## Methods

### Study approval

All experiments involving human subjects were conducted in accordance with the principles of the Declaration of Helsinki and were approved by the ethics committee of Kyoto University Hospital (protocol numbers: R0091, G0259, and G0457). Written informed consent was obtained from the legal guardians of the patients.

### Generation of iPSCs, iPS-MLs, and iPS-MPs

iPSCs from the patients with the *CDC42*^R186C^ variant were generated from peripheral blood mononuclear cells (PBMCs) (Pt.1) and fibroblasts (Pt.2) as described previously (31). Control iPSC lines were provided by the RIKEN BioResearch Center through the National Bio-Resource Project of the Ministry of Education, Culture, Sports, Science, and Technology, Japan. After confirming the quality of the iPSCs, including marker gene expression, absence of residual vector expression, and pluripotency, iPSC-MLs were established as previously described (8). In brief, floating hematopoietic cells that were differentiated into the monocytic lineage from iPSCs were collected on days 15–18. Cells were transfected with lentiviral constructs encoding BMI1, cMYC, and MDM2 in the CSII-EF-RfA vector kindly provided by Drs. Satoru Senju (Kumamoto University) and Hiroyuki Miyoshi (RIKEN Bio Resource Center),and cultured in StemPro-34 serum-free medium (Invitrogen) containing Glutamax (Gibco) in the presence of M-CSF (50 ng/mL, R&D Systems) and GM-CSF (50 ng/mL, R&D Systems). For macrophage differentiation, iPS-MLs were cultured in RPMI-1640 medium (Sigma-Aldrich) supplemented with 20% FBS and M-CSF (100 ng/mL) for 6 to 7 days. Adherent macrophages were detached with Accumax (Innovative Cell Technologies) and collected for subsequent experiments. The morphology and surface marker expression of mutant iPS-MLs and iPS-MPs were comparable to those of control counterparts.

### Single-base editing of CDC42 in iPSCs

Mutation correction (*CDC42*^R186C^ to *CDC42*^WT^) and mutagenesis (*CDC42*^WT^ to *CDC42*^R66G^) of the *CDC42* gene in iPSCs were performed by CRSPR-Cas9 adenosine base editing (32, 33). Two adenine base-editor expression vectors with the *EF-1a* promoter and puromycin–resistance gene, EF1a-NGG-ABEmax-puroR and EF1a-NG-ABEmax-puroR, were constructed based on pCMV_ABEmax (34), NG-ABEmax (35) (Addgene plasmid #112095, #124163, gifts from David Liu) and pHL-EF1a-SphcCas9-iP-A (36) (Addgene plasmid #60599, a gift from Akitsu Hotta). Single guide (sg) RNA target sites were “ctgcagctcttcttcggttc” for the correction of *CDC42*^R186C^ (c.556C>T), and “gatgacagattacgaccgctg” for the introduction of *CDC42*^R66G^ (c.A196A>G). sgRNA target oligos were incorporated into sgRNA plasmids: pFYF1320 EGFP Site#1 (37), a gift from Keith Joung (Addgene plasmid #47511; http://n2t.net/addgene:47511;RRID:Addgene_47511), according to a protocol from David Liu’s lab (https://benchling.com/protocols/E64XFSgM/sgrna-plasmid-cloning). sgRNA plasmid (5 µg) and EF1a-NGG/EF1a-NG-ABEmax-puroR (5 µg) were introduced into iPSCs by NEPA21 electroporator (Nepa Gene). Twenty-four hours after electroporation, puromycin (1 µg/mL) was added for an additional 24 h. Surviving colonies were picked up, and the genotypes were validated by Sanger sequencing.

### Generation of iPSCs carrying CDC42^WT^, CDC42^R186C^, CDC42^C188Y^, and CDC42^*192C*42^ by CRISPR-mediated homologous recombination

iPSC clones carrying *CDC42*^WT^, *CDC42*^R186C^, *CDC42*^C188Y^ and *CDC42*^*192C*24^ on otherwise identical genetic backgrounds were generated by CRISPR-mediated homologous recombination, as described previously (38). In brief, two sgRNA-Cas9 expression vectors (PX458, Addgene # 43138) targeting exon 7 and intron 6 of the *CDC42* gene and targeting vector with a floxed puromycin resistance cassette flanked by 2 kb homology arms were introduced into an iPSC clone derived from a healthy control for homologous recombination. The targeting vectors were constructed with PCR-amplified 3’ and 5’ homology arms, puromycin-resistance cassette, and backbone pENTR-DMD-Donor vector (Addgene, #60605) using HiFi DNA Assembly (New England Biolabs). To avoid repeated digestion after successful recombination, synonymous mutations were introduced into PAM sequences of the targeting vector by PCR-based mutagenesis. Each CDC42 variant was also introduced into the corresponding targeting vector by PCR-based mutagenesis. Two sgRNA-CRISPR/Cas9 vectors (2.5 µg each) and the targeting vector for each mutant (5 µg) were introduced into 1.0 × 10^6^ iPSCs using an NEPA 21 electroporator (Nepa Gene). Forty-eight to seventy-two hours after electroporation, 1 µg/mL puromycin was added. Surviving colonies were picked up and the genotypes were validated by Sanger sequencing. After genotype confirmation, the puromycin–resistance cassette was removed by introduction of the Cre recombinase expression vector.

### mRNA silencing

On day 3 of differentiation from iPS-MLs to iPS-MPs, cells were transfected with 60 ng Silencer Select predesigned small interfering RNA (siRNA; CDC42 siRNA ID: s2765/s55424, MEFV siRNA ID: s502555/s502557; Thermo Fisher Scientific) or Silencer Select Negative Control No. 1 siRNA (Thermo Fisher Scientific) using the Lipofectamine RNAiMAX transfection reagent (Thermo Fisher Scientific).

### *In vitro* stimulation of iPS-MLs and iPS-MPs

iPS-MLs or iPS-MPs were harvested as described above and seeded in 96-well plates at 5 × 10^4^ cells per well. When indicated, cells were pretreated with ABT-751(10 µM, Sigma-Aldrich), CA4P (10 µM, Sigma-Aldrich), MCC950 (10 µM, Sigma-Aldrich) and colchicine (100 ng/mL, Sigma-Aldrich) 30 minutes before LPS priming, or with 2-bromo-palmitate (20 µM, Sigma-Aldrich) and bryostatin (0.1 µM, Sigma-Aldrich) 24 h before harvest. For inflammasome activation, cells were primed with LPS (1 µg/mL) for 4 h and treated with UCN-01 (10 µM, Sigma-Aldrich), nigericin (10 µM, Sigma-Aldrich), or purified flagellin from S. typhimurium (InvivoGen) in DOTAP liposomes (Roche) (IC-FLA; 6 µg DOTAP/1 µg FLA-ST) for an additional 2 h, after which the supernatants were collected. For TcdA stimulation, TcdA (1 µg/mL, List Biological Laboratories) was added after 2 h of LPS priming (1 µg/mL), and supernatants were collected 4 h later. For AIM2 inflammasome activation, poly (dA; dT)/ Lyovec (1 µg/mL, InvivoGen) was added along with LPS priming (1 µg/mL), and supernatants were collected 4 h later. The IL-1β, IL-18 and IL-6 concentrations were measured using the Bio-Plex Pro Human Cytokine Assay (BioRad).

### Immunofluorescence staining and quantification of ASC aggregates

iPS-MPs were stimulated as in the cytokine secretion assay. After 2 h of stimulation with TcdA, the cells were attached to slides with a Cytospin 4 Cytocentrifuge (Thermo Fisher Scientific), fixed in 4% paraformaldehyde, and permeabilized with 0.1% Triton X-100. Cells were incubated with an anti-ASC antibody and then with an Alexa Fluor 594-labeled antibody against rabbit IgG. Nuclei were stained with Hoechst. The cells were examined by fluorescence microscopy on a BZ-X710 microscope (Keyence), and the BZ-X Analyzer software (Keyence) was used for quantitative analysis.

### Plasmids

The N-terminal FLAG-tagged WT human *CDC42* gene (isoform 1, NP_001782.1) and the puromycin– resistance gene separated by a T2A peptide were inserted into the pcDNA5/TO Mammalian Expression Vector by HiFi DNA Assembly to generate a plasmid encoding FLAG-CDC42-t2a-puromycin resistance. An N-terminal-AcGFP-fused CDC42 expression vector was constructed by incorporating PCR-amplified *CDC42* cDNA into the pAcGFP1-C1 Vector (Takara Bio). Each *CDC42* variant was generated by PCR-based mutagenesis using KOD plus (Toyobo). For co-expression assays with *CDC42* and *MEFV*, a C-terminally GFP-fused *MEFV* gene (isoform 1, NP_000234.1) and *CDC42* separated by T2A peptides were inserted into the pAcGFP1-C1 Vector to obtain plasmids encoding hMEFV-pAcGFP1-t2a-CDC42.

### Subcellular localization of CDC42

COS-1 cells were transfected with AcGFP-CDC42 using PEI MAX (Polysciences). Twenty-four hours after transfection, cells were fixed with 4% paraformaldehyde, permeabilized with 0.1% Triton X-100, and quenched with 50 mM NH4Cl. After blocking with 3% BSA in PBS, cells were incubated with primary antibodies followed by secondary antibodies conjugated to Alexa fluorophore and then mounted with ProLong Glass Antifade Mountant (P36982, Thermo Fisher Scientific). Nuclei were stained with DAPI (11034-56, Nacalai Tesque). Confocal microscopy was performed using LSM880 with Airyscan (Zeiss) with a 100 × 1.46 alpha-Plan-Apochromat oil immersion lens. Images were analyzed and processed with Zeiss ZEN 2.3 SP1 FP3 (black, 64 bit) (ver. 14.0.21.201) and Fiji (ver. 2.0.0-rc-69/1.52p).

### CDC42 transient expression for immunoprecipitation analysis

Plasmids encoding the FLAG-CDC42(WT/mutant)-t2a-puromycin resistance gene or empty vector were transfected into HEK293T cells using Trans-IT 293. Forty-eight hours after transfection, puromycin (1μg/mL) was added for an additional 24 h. The cells were then harvested and assayed for immunoprecipitation analysis.

### CDC42 pull down activation assay

The active, GTP-bound form of CDC42 was pulled down for analysis using the Cdc42 Pull-Down Activation Assay Kit (Cytoskeleton). FLAG-tagged protein pull-down assays were performed using DDDDK tagged protein purification kit (MBL). Samples were subjected to western blotting for further analysis.

### Acyl-biotin exchange assay for detecting palmitoylated CDC42

The acyl–biotin exchange assay was performed according to a protocol by Junmei Wan (39) with minor modifications. HEK293T cells were transfected with the N-terminal FLAG-tagged *CDC42* expression vector using TransIT 293 (Mirus Bio LLC) in 6-well culture plates. After 48 h of transfection, the cells were lysed and subjected to assays. In brief, cell lysates were treated with N-ethylmaleimide to block unmodified cysteine-residues on proteins. Palmitoylated cysteines on proteins are cleaved and unmasked by hydroxylamine (HAM) treatment, yielding free cysteines that were previously palmitoylated. Free cysteines were subsequently treated with thiol-reactive HPDP-biotin, and biotinylated proteins were pulled down with streptavidin-beads. Eluted proteins were analyzed by western blot. Every assay includes a non-HAM treated control to distinguish palmitoylated proteins from nonspecifically biotinylated proteins. A detailed protocol is available upon request.

### Western blotting

Protein samples were separated by SDS-PAGE, transferred to PVDF membrane, and probed with specific primary antibodies (Abs) followed by the horseradish peroxidase (HRP)-conjugated secondary antibodies. Specific bands were visualized by enhanced chemiluminescence using the Clarity Western ECL Substrate (Bio-Rad) and quantified by the ChemiDoc XRS1 System and Image Laboratory software (Bio-Rad). Mouse monoclonal anti-CDC42 (Cytoskeleton, 1:1000), mouse polyclonal GAPDH (FUJIFILM Wako, 1:4000), rabbit polyclonal anti-human pyrin (AdipoGen, 1:2000), mouse monoclonal anti Rho-GDI (Santa-Cruz Biotechnology, 1:500) and mouse monoclonal anti-FLAG M2 Ab (Sigma, 1:1000), anti-ERK1/2 (Cell Signaling Technology, 1:1000), anti-p-ERK1/2 (Cell Signaling, 1:1000), anti-p65 (Cell Signaling, 1:1000), anti-p-p65 (Cell Signaling, 1:1000), anti-JNK (Cell Signaling, 1:1000), anti-p-JNK (Cell Signaling, 1:1000), anti-p38 (Cell Signaling, 1:1000), anti-p-p38 (Cell Signaling, 1:1000) served as primary Abs for western blotting. HRP-conjugated anti-mouse/rabbit IgG antibodies (Jackson ImmunoResearch Laboratories, 1:4000) were used as secondary antibodies.

### Nucleofection and Flow Cytometry

THP-1 nucleofection and cell death assays were performed as described previously (40, 41) with some modifications. THP-1 cells (1 × 10^6^) were transfected with 500 ng plasmid encoding pAcGFP-fused CDC42 variants or *hMEFV*-pAcGFP1-t2a-*CDC42* using 4D-Nucleofector and the SG Cell Line 4D-Nucleofector X Kit (Lonza). Immediately after nucleofection, 10 ng/mL phorbol 12-myristate 13-acetate (FUJIFILM Wako) was added, and the cells were cultured overnight. The next day, the cells were stained with LIVE/DEAD Fixable Violet Dead Cell Stain Kit (Thermo Fisher Scientific) as indicated. Cells were analyzed using a FACSVerse flow cytometer (BD; Becton and Dickinson Bioscience) and FlowJo software (BD). Cell death was calculated as the percentage of the size-gated cells that were BV421-high.

### NK cell cytolytic assay

PBMCs were stimulated and expanded in NK MACS^®^ Medium (Miltenyi Biotec) for more than 3 weeks. These cells were co-cultured for 6 h with K562 cells in 96-well microtiter plates as targets, and cytolytic activity was determined in an LDH release assay (Non-Radioactive Cytotoxicity Assay; CytoTox 96 Assay, Promega). Background spontaneous and maximal LDH release were determined by culturing target cells in medium alone or in medium containing Triton X-100.

### Degranulation assays

To quantify lysosome exocytosis by NK cells, 2 × 10^5^ PBMCs stimulated for 48 h with human recombinant interleukin-2 (rIL-2, 50 U/mL), were cultured with or without 2×10^5^ K562 cells and incubated for 2 h. Lysosomal degranulation of cytolytic T cells (CTLs) were evaluated using alloantigen-specific CD8+ CTL lines as described previously (42, 43). Briefly, PBMCs obtained from patients and healthy individuals were co-cultured with a mitomycin C (MMC)-treated B-lymphoblastoid cell line (KI-LCL) established from an HLA-mismatched individual. Six days later, CD8+ T lymphocytes were isolated using immunomagnetic beads (MACS beads; Miltenyi Biotec). The cells were cultured in RPMI 1640 medium supplemented with 10% human serum and 10 IU/mL human rIL-2, and stimulated weekly with MMC-treated KI-LCL cells for more than 3 weeks. These lymphocytes (2 × 10^5^ in 96-well round bottom plates) were mixed with an equal number of P815 cells in the presence of 0.5 µg/mL anti-CD3 mAb (OKT3) for 2 h. Cells were resuspended in PBS supplemented with 2% fetal calf serum and 2 mM EDTA; stained with anti-CD3, anti-CD8, anti-CD56, and anti-CD107a mAbs; and analyzed by flow cytometry.

### Statistical analysis

Statistical significance was determined by one-way ANOVA with Tukey’s test, Student’s t-test, or ratio paired t-test as indicated in the figure legends. All statistical analyses were performed using GraphPad Prism (ver. 8.2.1; GraphPad Software).

## Acknowledgments

The authors are grateful to the patients and their families. This work was supported by JSPS KAKENHI (Grant Numbers: 21K07795, 20H03202, and 19H00974), AMED (Grant Numbers: JP20ek0109387, JP21bm0104001, JP21bm0804004, and JP19bm0804001), AMED-PRIME (Grant Number: 17939604), and the “Research on Measures for Intractable Diseases” project from the Japanese Ministry of Health, Labor, and Welfare.

## Authorship

Contributions: R. N., T. Taguchi, Y. S., and T. Y. designed the research; M. N-I. performed genetic manipulation of iPSCs, cell culture and stimulation assays, western blotting, and pull down assays; K. M. performed immunocytochemical analyses; O. O. performed genetic analysis; Y. Kawasaki., M. O., and M. K. S. generated iPSCs and iPS-MLs; Y. H., H. N., T. Tanaka, H. S., K. K. assisted the research; M. N-I., Y. H., H. N., T. Tanaka, H. S., K. I., Y. Katata, S. O., T. W., R. N., and T. Y. attended to the patients; M. N-I., K. M., Y. H., T. Tanaka, E. H., K. I., S. K., J. T., R. N., T. Taguchi., Y. S., and T. Y. analyzed the data and discussed the results; M. N-I., K. M., Y. H., and T. Y. wrote the paper.

