## Supplemental Materials for "Aberrant localization of CDC42 C-terminal variants to the Golgi apparatus drives pyrin inflammasome-dependent autoinflammation"

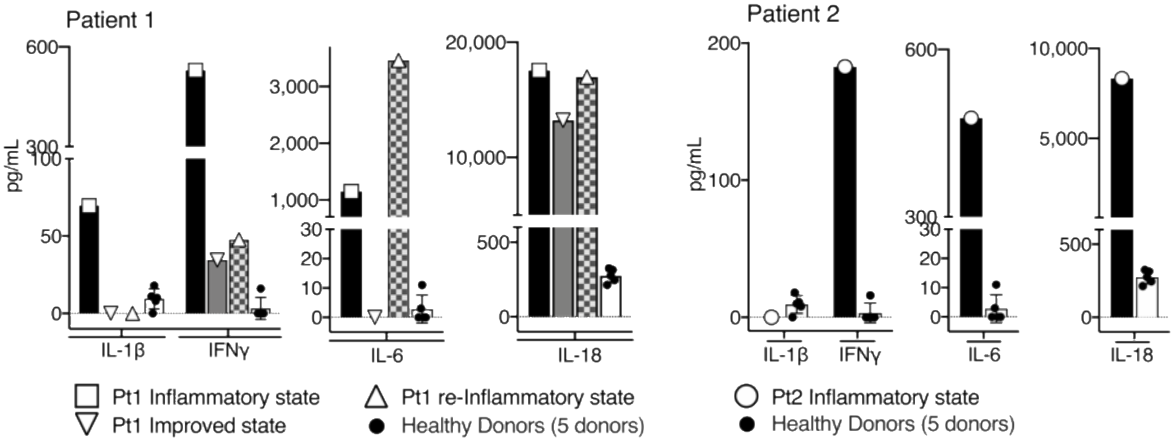


**Supplementary Figure 1. Elevated levels of IL-18 in sera of patients with CDC42$^{R186C}$.**

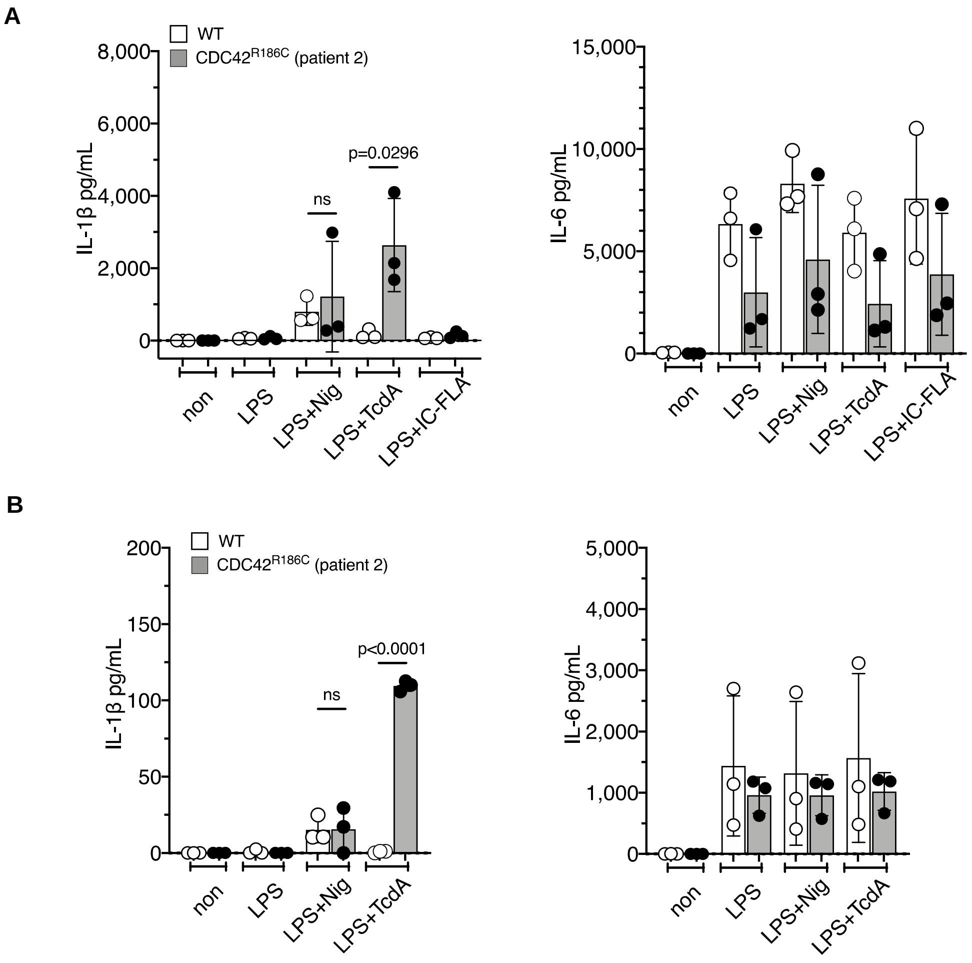


**Supplemental Figure 2. Response to TcdA stimulation is also increased in iPS-MPs and iPS-MLs derived from Pt.2.** IL-1β and IL-6 release in response to various inflammasome stimuli from iPS-derived (**A**) MPs and (**B**) MLs established from Pt. 2 and healthy controls. Data are representative of three independent experiments using three independent iPS clones. Statistical significance was determined by one-way ANOVA with Tukey's multiple comparison.

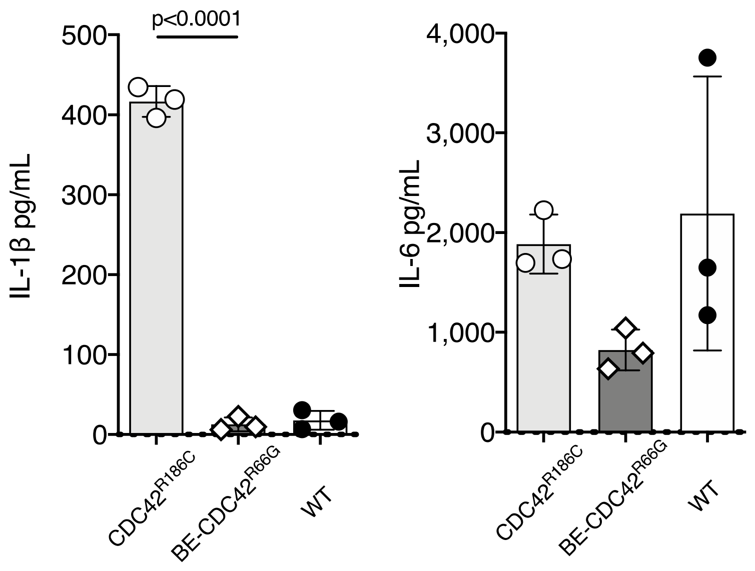


**Supplemental Figure3. IL-1β production by iPS-MLs harboring CDC42$^{R66G}$ is comparable to that of CDC42$^{WT}$ cells.** iPSCs carrying CDC42$^{R66G}$ were generated by manipulating WT-iPSCs by single-base editing and iPS-MPs were differentiated. Cells were stimulated with LPS+TcdA and the production of IL-1β and IL-6 was evaluated. Representative results from three independent experiments using three independent clones are shown. Statistical significance was determined by one-way ANOVA with Tukey's multiple comparison.

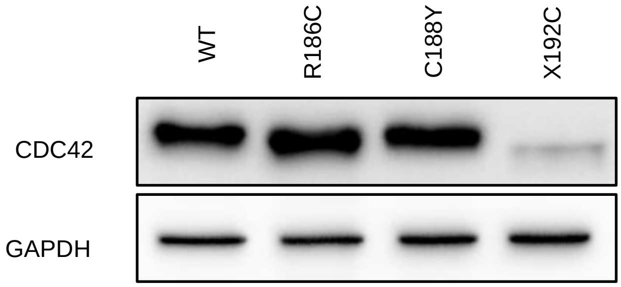


**Supplemental Figure 4. CDC42$^{*192C*24}$ is less stable than other CDC42 C-terminal variants.** CDC42 C-terminal variants were transiently transfected into HEK293T cells, and their expression levels were evaluated by Western blot analysis.

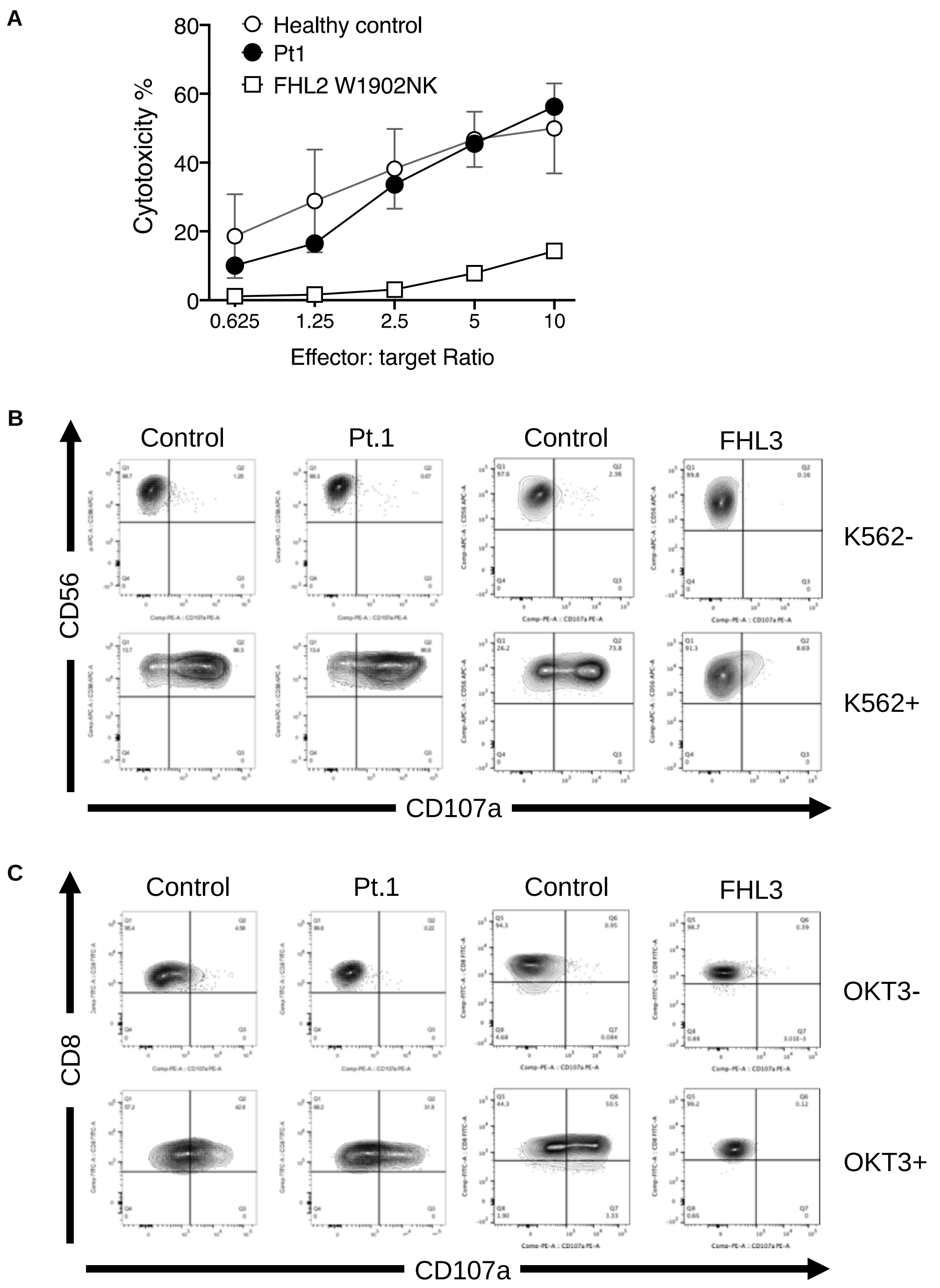


**Supplemental Figure 5. Cytolytic and degranulation activity of NK cells and CTLs from Pt.1.** (**A**) Cytolytic activity of activated NK cells derived from healthy controls, Pt.1, and a FHL2 patient. Degranulation of activated (**B**) NK cells and (**C**) CTLs derived from healthy controls, Pt.1, and a FHL3 patient. NK, Natural Killer. FHL, familial hemophagocytic lymphohistiocytosis.

### Case Report

Patient 1 was a first male child born at term after a noneventful pregnancy to healthy, non-consanguineous Japanese parents. Starting on the first day of life, he presented with fever, erythema, diarrhea, and hepatosplenomegaly. C-reactive protein (CRP) rose to 182 mg/L, and pancytopenia (neutrophils 600/μL, hemoglobin 10.3g/dL, platelets 14.9 × 109/L) was noted. No dysmorphic features were observed, and the patient's platelet size was within normal range. Extensive microbiological screening returned negative results. Fasting and intravenous administration of high-dose immunoglobulin were partially effective, but their effects were transient, and the patient's condition gradually worsened. Prednisolone (2 mg/kg/day) was started, and the patient's condition stabilized with recovery of laboratory parameters. CRP became negative, and the count of neutrophils and platelets reached 1,650/μL and 22.5 × 109/L, respectively. However, fever and erythema recurred after gradual tapering of prednisolone. CRP level elevated and pancytopenia worsened, requiring recurrent transfusion of red blood cells and platelets. Bone marrow biopsy revealed hypocellular marrow without remarkable hemophagocytosis. Reescalation of prednisolone could not suppress inflammation. Etanercept was introduced, but no obvious effect was observed, and the patient died at 4.5 months of age due to overwhelming inflammation.

Patient 2 was a first male child born with body weight of 2194 g at 36 weeks of gestation to healthy nonconsanguineous Japanese parents. On day 6, he presented with fever, erythema multiforme, diarrhea, hepatosplenomegaly, and cholestasis. CRP rose to 48.9 mg/L, and pancytopenia (neutrophils 860/μL, hemoglobin 12.2g/dL, platelets 72.0 × 109/L) was noted. The patient's platelet size was within the normal range, and he exhibited no dysmorphic features. Extensive screens for infection were all negative, and no response was observed to antibacterial and antiviral treatments. Bone marrow examination revealed hypoplastic marrow with scattered foam cells. Dexamethasone therapy was initially effective for the control of symptoms and laboratory parameters. However, CRP level increased to 380 mg/L after gradual tapering of dexamethasone. Subsequent steroid therapy with prednisolone and methylprednisolone pulse therapy achieved limited control of intractable inflammation and pancytopenia. Finally, we treated him with plasma exchange therapy. However, the patient died at 2.5 months of age due to acute respiratory distress syndrome caused by durable and worsening systemic inflammation.

Whole-exome sequencing was performed, and a de novo heterozygous c.556C>T variant in CDC42 was identified in both patients.
